# Acetate drives ovarian cancer quiescence via ACSS2-mediated acetyl-CoA production

**DOI:** 10.1101/2024.07.12.603313

**Authors:** Allison C. Sharrow, Emily Megill, Amanda J. Chen, Afifa Farooqi, Stacy McGonigal, Nadine Hempel, Nathaniel W. Snyder, Ronald J. Buckanovich, Katherine M. Aird

**Author notes:** **Correspondence:** Katherine M. Aird Associate Professor Department of Pharmacology & Chemical Biology UPMC Hillman Cancer Center University of Pittsburgh School of Medicine 5051 Centre Ave. Office: 2041; Lab: 2050 Pittsburgh, PA 15213 and Ronald J. Buckanovich Professor Division of Gynecologic Oncology, Department of Obstetrics and Gynecology Department of Medicine UPMC Hillman Cancer Center University of Pittsburgh School of Medicine and Magee-Womens Research Institute 204 Craft Avenue Pittsburgh, PA 15213.

## Abstract

Quiescence is a reversible cell cycle exit traditionally thought to be associated with a metabolically inactive state. Recent work in muscle cells indicates that metabolic reprogramming is associated with quiescence. Whether metabolic changes occur in cancer to drive quiescence is unclear. Using a multi-omics approach, we found that the metabolic enzyme ACSS2, which converts acetate into acetyl-CoA, is both highly upregulated in quiescent ovarian cancer cells and required for their survival. Indeed, quiescent ovarian cancer cells have increased levels of acetate-derived acetyl-CoA, confirming increased ACSS2 activity in these cells. Furthermore, either inducing ACSS2 expression or supplementing cells with acetate was sufficient to induce a reversible quiescent cell cycle exit. RNA-Seq of acetate treated cells confirmed negative enrichment in multiple cell cycle pathways as well as enrichment of genes in a published G0 gene signature. Finally, analysis of patient data showed that ACSS2 expression is upregulated in tumor cells from ascites, which are thought to be more quiescent, compared to matched primary tumors. Additionally, high *ACSS2* expression is associated with platinum resistance and worse outcomes. Together, this study points to a previously unrecognized ACSS2-mediated metabolic reprogramming that drives quiescence in ovarian cancer. As chemotherapies to treat ovarian cancer, such as platinum, have increased efficacy in highly proliferative cells, our data give rise to the intriguing question that metabolically-driven quiescence may affect therapeutic response.

## Introduction

Quiescence is defined as a reversible cell cycle exit at the G_0_ stage, which is promoted by the cyclin dependent kinase inhibitors p21 (encoded by *CDKN1A*), p27 (*CDKN1B*), and p57 (*CDKN1C*) [1-5]. Quiescent cells have generally been considered to be metabolically inactive because of fewer biosynthetic requirements [6]. However, recent data indicate that quiescent cells are metabolically active but with distinct metabolic profiles from proliferative cells [7, 8]. Most of these studies have been performed in the context of normal quiescent stem cells, and how metabolism is altered in cancer cell quiescence is less clear.

Tubo-ovarian high grade serous carcinoma (HGSC) is the most common and most deadly form of ovarian cancer [9]. Most patients with HGSC present with stage III-IV disease, and surgery combined with platinum and taxane based chemotherapy remains the primary treatment. Unfortunately, despite initial response to therapy, 70-90% of patients with stage III-IV HGSC will experience disease recurrence after first-line therapy [10, 11]. Once a patient’s ovarian cancer returns, her median survival time is only ∼18 months [10, 11]. Chemotherapies, such as the platinum-based therapies used to treat HGSC, target rapidly proliferating cells, and quiescent cells are therefore inherently more chemoresistant. Indeed, after the cessation of chemotherapy, these cells can re-enter the cell cycle to drive relapse [5, 12, 13]. Our prior studies found that platinum induces quiescence in ovarian cancer cells, promoting resistance in both cell-intrinsic and cell-extrinsic manners [14, 15]. How other cell-intrinsic or microenvironmental pathways induce quiescence in ovarian cancer remains underexplored.

In this study, we sought to identify metabolic changes that influence HGSC quiescence. Using two unbiased approaches, we identified acyl-CoA synthetase short chain family member 2 (ACSS2) to be both upregulated in quiescent cells and important for their survival. ACSS2 converts acetate to acetyl-CoA. Consistent with upregulation of ACSS2, we found increased production of acetyl-CoA from acetate in multiple HGSC quiescent models. Similarly, both overexpression of ACSS2 or supplementing cells with acetate induced HGSC cell quiescence. Linking ACSS2 to a role in human cancer, we found that increased expression of *ACSS2* mRNA, or the quiescence regulatory gene *CDKN1C* mRNA, correlates with a worse patient outcomes. Finally, we found that ACSS2 expression is greater in cancer cells from ascites (which are generally more quiescent) than solid tumor associated cancer cells. Together, our data demonstrate that acetate-derived acetyl-CoA drive HGSC quiescence and suggest that this axis is a potential therapeutic target to overcome chemoresistance.

## Materials and Methods

### Cell culture and metabolites

Ovcar8 and Kuramochi cells were a gift from Dr. Benjamin Bitler (University of Colorado). Cells were cultured in RPMI-1640 (Corning; Corning, NY, USA) with 5% fetal bovine serum (VWR Seradigm), 100 U/mL penicillin (Gibco), and 100 U/mL streptomycin (Gibco) HEK293FT cells were used for lentiviral packaging and were cultured in DMEM (Corning, cat#10-013-CV) supplemented with 10% FBS according to ATCC. For acetate treatment, cells were supplemented with sodium acetate (Sigma cat#S2889), sodium citrate (C7254), BSA-oleate monosaturated fat (Cayman Chemical, cat#295575), or sodium butyrate (Sigma, cat#B5887) for 72h unless otherwise specified. All cell lines were tested monthly for mycoplasma as described [16].

### Induction of quiescence

Cells were induced to quiesce using two approaches: serum starvation and culturing in non-adherent conditions. Serum starvation-induced quiescence: Ovcar8 cells were cultured without FBS for 48h (the timepoint at which we observed decreased proliferation without decreased viability). Kuramochi cells were cultured without FBS for 96h. Non-adherence-induced quiescence: Cells were seeded in ultra-low attachment plates (Corning, cat#3471) and cultured for 48h.

### Plasmids and lentiviral production

Human pooled metabolic CRISPR KO library (Addgene #110066), pLV-mCherry-hCdt1(1-100)ΔCy (Addgene #193759), and pCDH-EF1-mVenus-p27K^−^ (Addgene #176651) were obtained from Addgene. The human ACSS2 lentiviral expression plasmid was synthesized by Twist Biosciences (San Francisco, CA, USA). Lentivirus was packaged using the ViraPower Kit (Invitrogen, cat#K497500) following the manufacturer’s instructions. Cells were infected with corresponding vectors for 16h and selected for 3 days with 1µg/mL of puromycin.

### RNA-Seq

Total RNA was extracted from cells with Trizol (Ambion, cat#15596018) and DNase treated, cleaned and concentrated using Zymo columns (Zymo Research, cat#R1013) following the manufacturer’s instructions. RNA integrity number (RIN) was measured using BioAnalyzer (Agilent Technologies) RNA 6000 Nano Kit to confirm RIN above 7 for each sample. The cDNA libraries, next generation sequencing, and bioinformatics analysis was performed by Novogene. Raw and processed RNA-Seq data can be found on GEO (GSE271035 and GSE271357). Processed data can be found in **Table S1 and S2**.

### CRISPR knockout screen

Ovcar8 cells were transduced with lentivirus containing the pooled metabolic CRISPR KO library at a multiplicity of infection (MOI) of 0.2 to ensure single gene targeting per cell. Selection was conducted with 1µg/mL of puromycin for 3 days. Cells were passed every 2 days and the whole population was seeded to maintain 160X library coverage throughout. After selection, cells were harvested for genomic DNA extraction using the Zymo Research kit (cat# D4069). sgRNA inserts were PCR amplified using Ex Taq DNA Polymerase (Takara, cat#RR001A) from sufficient genome equivalents of DNA to achieve an average coverage of >200x of the sgRNA library. gRNA was amplified using primers: universal forward primer- AATGATACGGCGACCACCGAGATCTACAC- CGACTCGGTGCCACTTTT; reverse primer with index (control)- CAAGCAGAAGACGG- CATACGAGATCTATGGACCTTTCTTGGGTAGTTTGCAGTTTT; reverse primer with index (serum starved)- CAAGCAGAAGACGGCATACGAGATCCGACAGAATTTCTT-GGGTAGTTTGCAGTTTT. Pooled PCR amplicons were then sequenced. MAGeCK was used as the bioinformatics pipeline to analyze negatively and positively enriched genes [17]. Briefly, read counts from the samples were median-normalized to adjust for library size effect and read count distribution and mean-variance modeling was used to capture the relationship of mean and variance. Next, negative binomial (NB) model was used to assess whether there are significant differences in sgRNA abundance between serum starved and control groups similarly to as the method used for differential expression analysis in bulk RNA-Sequencing [18]. P-values are then calculated from the NB model and used to rank sgRNAs using an α-RRA algorithm [19] to identify positively or negatively selected genes. α-RRA operates under the assumption that if a gene has no impact on cell survival, target it using a sgRNA should be evenly distributed in the ranked list of all sgRNAs. Therefore, the algorithm ranks genes by comparing the skew in rankings to a uniform null model, prioritizing genes with consistently higher or lower-than-expected sgRNA rankings. The statistical significance of the skew is determined through permutation as described in [17]. **Table S3** contains the gene names (Gene), Pathway, p-values (p-value), log_2_-fold change (serum starved vs. control, lfc) and scores (score) for the CRISPR KO screen. While intergenic controls were included in both libraries and use for the bioinformatics analysis, they have been removed from the graph and table.

### Western Blotting

Cells lysates were collected in 1X sample buffer (2% SDS, 10% glycerol, 0.01% bromophenol blue, 62.5mM Tris, pH 6.8, 0.1M DTT) and boiled to 95°C for 10 min. Protein concentration was determined using the Bradford assay (Bio-Rad, cat#5000006). An equal amount of total protein was resolved using SDS-PAGE gels and transferred to nitrocellulose membranes (Cytiva, cat#10600001) at 110mA for 2h at 4°C. Membranes were blocked with 5% nonfat milk in TBS containing 0.1% Tween-20 (TBS-T) for 1 h at room temperature. Membranes were incubated overnight at 4°C in primary antibodies diluted in 4% BSA/TBS + 0.025% sodium azide. Membranes were washed 4 times in TBS-T for 5 min at room temperature after which they were incubated with HRP-conjugated secondary antibodies for 1 h at room temperature. After washing 4 times in TBS-T for 5 min at room temperature, proteins were visualized on film after incubation with SuperSignal West Pico PLUS Chemiluminescent Substrate (Thermo Scientific, cat#34580). Primary antibodies: anti-AceCS1 (1:1000, clone: D19C6, Cell Signaling Technology) and anti-β-actin (1:10,000, clone: AC-15, Millipore Sigma). Secondary antibodies: anti-rabbit IgG-HRP (1:5000, cat#7074, Cell Signaling Technology) and anti-mouse IgG-HRP (1:5000, cat#7076, Cell Signaling Technology).

### Proliferation and Viability Assays

Cells were transduced with Nuclight Green Lentivirus (Sartorius) and selected for cells expressing nuclear-restricted GFP2. Longitudinal cell death was measured by including 250 nM Cytotox Red (Sartorius) in cultures. GFP2-expressing cells were plated into 96-well plates at a density of 3.4×10^3^ cells/well for Ovcar8 and 6×10^3^ cells/well for Kuramochi. For serum starvation, cells were permitted to adhere overnight, then medium was changed on both serum-starved and control wells, and cell numbers were monitored in an Incucyte S3 live-cell analysis system (Sartorius).

### Flow cytometry analysis

Cells expressing both pLV-mCherry-hCdt1(1-100)ΔCy and pCDH-EF1-mVenus-p27K^−^ were stimulated to quiesce, and cell cycle status was analyzed using a LSR Fortessa flow cytometer (BD Biosciences). FlowJo was used to quantify the double positive, quiescent population as described previously [20].

### Gene Set Enrichment Analysis

Gene Cluster Text files (GTC), as well as Categorical Class files (CLS) were generated independently for RNA-Seq datasets following the Gene Set Enrichment Analysis (GSEA) documentation indications (http://software.broadinstitute.org/gsea/index.jsp). GTC and CLS files were used to run independent GSEA analysis (javaGSEA desktop application). GSEA for Hallmarks, KEGG, and WikiPathways were run independently under the following parameters: 1000 permutations, weighted enrichment analysis, signal to noise metric for ranking genes, and “meandiv” normalization mode. Following GSEA documentation indications, terms with a q-value < 0.25 were considered significant (**Table S4**).

### DepMap data

For correlation of gene expression, DepMap Expression Public 24Q2 was downloaded for ovary/fallopian tube cell lines (June 2024). For GSEA analysis based on ACSS2 expression, the lower and upper quartiles were used.

### Acetyl-CoA Quantification and Tracing

For steady state abundance, cells were scraped into 10% trichloroacetic acid. For isotopologue tracing, cells were transferred to medium containing 1 mM sodium acetate (1,2-¹³C₂) (Cambridge Isotope Lab; Tewksbury, MA, USA) or 5 mM D-glucose-^13^C_6_ (Millipore Sigma) for the final 2 hours of treatment then scraped or centrifuged and resuspended into 10% trichloroacetic acid. Acyl-CoAs were measured by liquid chromatography-high resolution mass spectrometry for both quantification and isotope tracing as previously described in detail [21]. Briefly, for quantification, samples spiked with 0.1 mL of ^13^C_3_^15^N_1_-acyl-CoA internal standard prepared as previously published [22] and 0.9 mL of 10% (w/v) trichloroacetic acid in water. Calibration curves were prepared from commercially available acyl-CoA standards. Calibration curve samples were also subjected to sonication and extraction. Samples were homogenized using a probe tip sonicator for 0.5 second pulses 30 times then samples were centrifuged at 17,000 x g for 10 min at 4°C. Supernatant was purified by solid phase extraction (SPE) cartridges (Oasis HLB 10 mg, Waters) conditioned with 1 mL of methanol then 1 mL of water. Acid-extracted clairified supernatants were loaded onto the cartridges and washed with 1 mL of water. Acyl-CoAs were eluted with 1 mL of 25 mM ammonium acetate in methanol and evaporated to dryness under nitrogen gas. Samples were resuspended in 50 µL of 5% (w/v) 5-Sulfosalicyilic acid and 10 µL injections were analyzed on an Ultimate 3000 UHPLC using a Waters HSS T3 2.1×100mm 3.5 µm column coupled to a Q Exactive Plus. Data was analyzed using Tracefinder 5.1 (Thermo) and isotopic normalization via FluxFix [23]. The analyst was blinded to sample identity during processing and quantification.

### Patient survival and chemoresistance data

Data for HGSC patient overall survival were downloaded from KMplot [24]. Analysis was restricted to “Serous” histology and “Contains Platin” chemotherapy. Progression-free survival based on upper and lower quartile. Overall survival based on median *ACSS2* expression. Chemoresistance data was generated using cBioportal Ovarian Serous Cystadenocarcinoma (TCGA, Nature 2011) with *ACSS2* mRNA expression: Low: <-0.4; High: >0.4.

### Immunocytochemistry

Paired samples of human ovarian cancer tumor and ascites were centrifuged onto slides and fixed in 4% paraformaldehyde. Cells were permeabilized with 0.2% triton X-100, blocked with 3% BSA in PBS, probed for ACSS2 with anti-AceCS1 (1:50 dilution in 3% BSA in PBS, clone: D19C6, Cell Signaling Technology), detected with FITC-conjugated AffiniPure Donkey Anti-Rabbit IgG (1:1000 dilution in 3% BSA in PBS; Jackson ImmunoResearch), and counterstained with DAPI. Images of at least 50 cells per condition were captured using a Nikon TS100 fluorescent microscope. Positive staining for ACSS2 was quantified using thresholding for ACSS2 fluorescent staining and DAPI in FIJI.

### Quantification and statistical analysis

GraphPad Prism (version 9.0) was used to perform statistical analysis. Point estimates with standard deviations or standard errors were reported, as indicated, and the appropriate statistical test was performed using all observed experimental data. All statistical tests performed were two-sided and p-values<0.05 were considered statistically significant.

## Results

### Multi-omics analysis identifies ACSS2 as a candidate regulator of quiescence in HGSC

To understand metabolic changes that occur in HGSC quiescence, we performed both RNA-Seq and a metabolic CRISPR KO screen on serum-starved Ovcar8 cells (**Fig. 1A-B and Table S1-S2**). Serum starvation is a classical model of quiescence, and quiescence induction in Ovcar8 cells was confirmed through transiently reduced proliferation and a modified FUCCI reporter that quantifies G0 through dual CDT1/p27 staining (**Fig. S1A-C**) [25]. Comparing the results of the two screens identified 6 hits that are both increased in expression and required for quiescence cell survival (**Fig. 1C**). In particular, *ACSS2* was markedly upregulated in quiescent cells (**Fig. 1D**), and data from DepMap demonstrates that *ACSS2* expression positively correlates with expression of the quiescence marker *CDKN1C* (encodes for p57) in ovarian cancer cell lines (**Fig. S1D**). We confirmed quiescence-induced upregulation of ACSS2 in multiple HGSC cell lines using both serum starvation and cells cultured in non-adherent conditions, another well-known model of quiescence induction (**Fig. 1E, Fig. S1A-C, and Fig. S1E-H**) [26, 27]. Together, these data demonstrate that ACSS2 expression is increased in quiescent cells.

**Figure 1.**
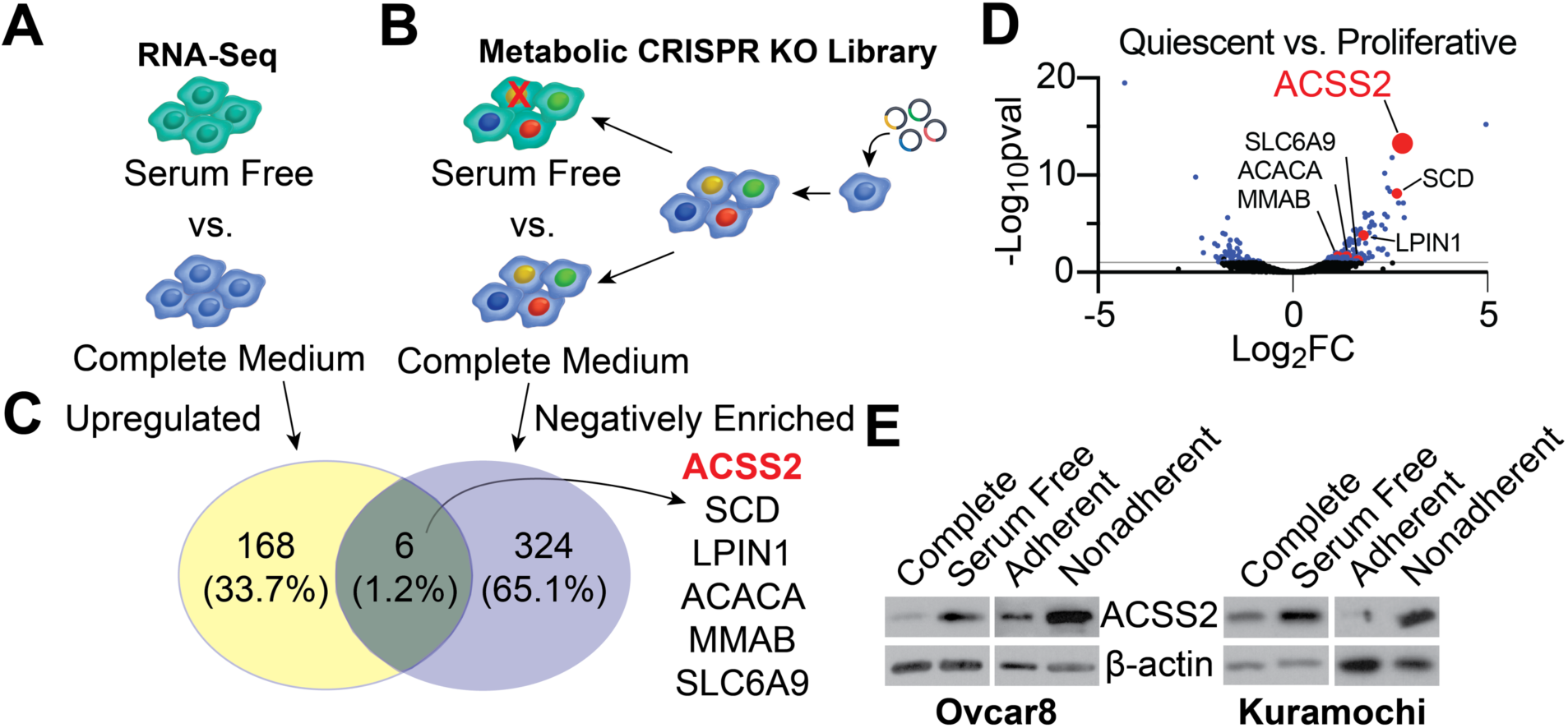
Multi-omics analysis identifies ACSS2 as a candidate regulator of quiescence. **(A-B)** Schematic representation of RNA-Seq **(A)** and CRISPR KO screen **(B)** of serum starved (serum free) Ovcar8 cells. **(C)** Cross-comparison of RNA-seq and CRISPR KO screen results (cutoffs for RNA-seq: fold-change ≥ 2, FDR ≤ 0.1; Cutoff for CRISPR screen: pval ≤ 0.1). (**D)** Volcano plot of RNA-seq of serum starved Ovcar8 results showing differential expression of common genes from **(C)**. **(E)** ACSS2 protein expression was determined by western blotting in the indicated cells. β-actin was used as a loading control. Western blots shown are representative data from at least 2 independent experiments in each cell line and condition.

### Acetate-derived acetyl-CoA is upregulated in quiescent HGSC

ACSS2 catalyzes the conversion of acetate to acetyl-CoA (**Fig. 2A**) [27, 28]. To determine whether the observed increase in ACSS2 corresponds to increased acetyl-CoA, we first performed steady state metabolomics on serum starved quiescent cells. Consistent with the upregulation of ACCS2 (**Fig. 1**), we observed an increase in acetyl-CoA abundance in quiescent Ovcar8 cells compared to proliferative controls (**Fig. 2B**). We next performed ^13^C-acetate isotopologue tracing to determine whether the increase in acetyl-CoA in quiescent cells is derived from acetate (**Fig. 2A**). Indeed, we observed an increase in acetate-derived acetyl-CoA in HGSC cells induced to quiesce through serum starvation or non-adherence (**Fig. 2C and Fig. S2A**). Glucose-derived acetyl-CoA was decreased in quiescent cells, suggesting that the increase in acetyl-CoA abundance is due to ACSS2 and not PDH or ACLY (**Fig. 2D-E and S2B**). Further supporting this conclusion, we did not observe a correlation between *CDKN1C* and *ACLY* or any of the PDH complex genes in the DepMap database (**Fig. S2C**). Together, these functional metabolic data indicate that acetate-derived acetyl-CoA is increased in quiescent HGSC cells.

**Figure 2.**
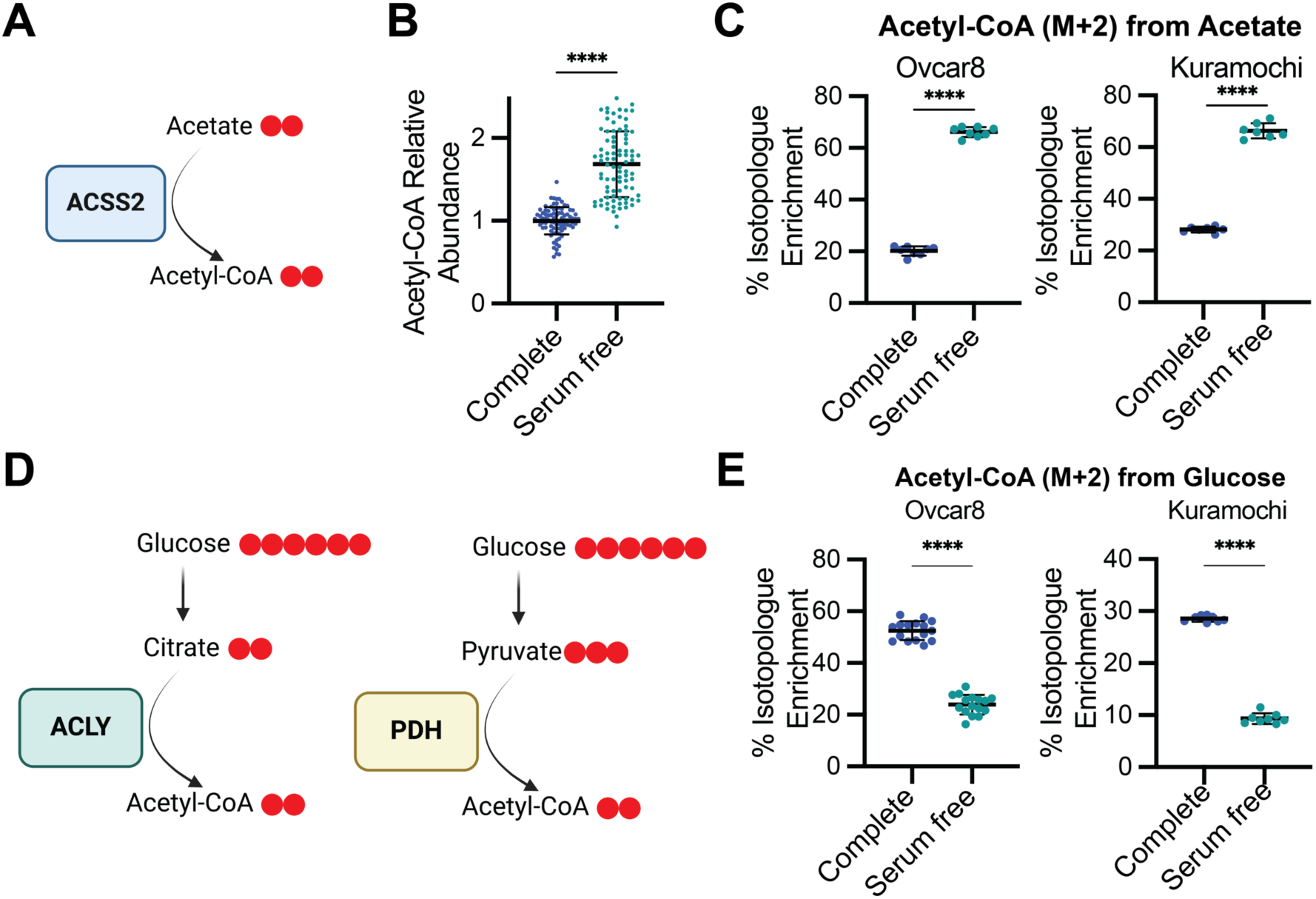
Acetate-derived acetyl-CoA abundance is increased in quiescent cells. **(A)** Schematic of acetate-mediated acetyl-CoA generation via ACSS2. Red dots indicate labeled carbons for isotopologue tracing. **(B)** Steady state acetyl-CoA abundance in Ovcar8 cultured in complete or serum free media was assessed by LC-MS. Pooled data from 6 independent experiments is shown (n=8). **(C)** Acetyl-CoA M+2 percent isotopologue enrichment from labeled acetate in the indicated cells. Representative data from at least 2 independent experiments in each cell line and condition. **(D)** Schematic of glucose-derived acetyl-CoA via ACLY and PDH. Red dots indicate labeled carbons for isotopologue tracing. **(E)** Acetyl-CoA M+2 percent isotopologue enrichment from labeled glucose in the indicated cells. Representative data from at least 2 independent experiments in each cell line and condition. Graphs represent mean ± SD. ****p<0.001

### Acetate and ACSS2 drive quiescence in HGSC models

We next sought to determine whether the observed upregulation of ACSS2 and increase in acetate-derived acetyl-CoA are a cause or consequence of quiescence. Consistent with a driver effect, ACSS2 overexpression reduced cell numbers without increasing cell death (**Fig. 3A-B and S3A**). To demonstrate the contribution of acetate to this phenotype, we supplemented cells with different doses of acetate. Indeed, acetate supplementation demonstrated a dose-dependent decrease in cell proliferation that did not markedly affect cell viability (**Fig. 3C-D and S3B**). We noted that Kuramochi cells required higher acetate concentrations to achieve a similar decrease in proliferation to Ovcar8, which may be due to cell line variability or differences in acetate uptake. Removal of acetate reversed the proliferative arrest (**Fig. 3E**), demonstrating a reversible effect on the cell cycle. The observed decrease in proliferation was acetate-specific, as supplementing with alternative acetyl-CoA sources failed to consistently decrease proliferation (**Fig. S3C**), although butyrate did decrease proliferation in one of the two cell lines. We confirmed that acetate-supplemented cells were enriched in the G_0_ phase (**Fig. 3F**), demonstrating induction of quiescence. Finally, we performed RNA-Seq on acetate supplemented cells and found the multiple cell cycle gene signatures were significantly downregulated (**Fig. 4A** - signatures in red and **Table S4**). Interestingly, a variety of metabolic gene signatures that are associated with proliferation were also decreased (**Fig. 4A**-signatures in blue). Moreover, there is a significant G0 signature [29] in acetate treated cells (**Fig. 4B and Table S5**; Downregulated G0 genes: NES= -1.74, FDR<0.001; Upregulated G0 genes: NES= 1.16, FDR=0.244). Finally, using data from DepMap, we also found a significant G0 signature in *ACSS2*-high versus *ACSS2*-low ovarian cancer cells (Downregulated G0 genes: NES= -2.19, FDR<0.001; Upregulated G0 genes: NES=1.24, FDR=0.174). These data demonstrate that ACSS2 and acetate are sufficient to drive quiescence induction.

**Figure 3:**
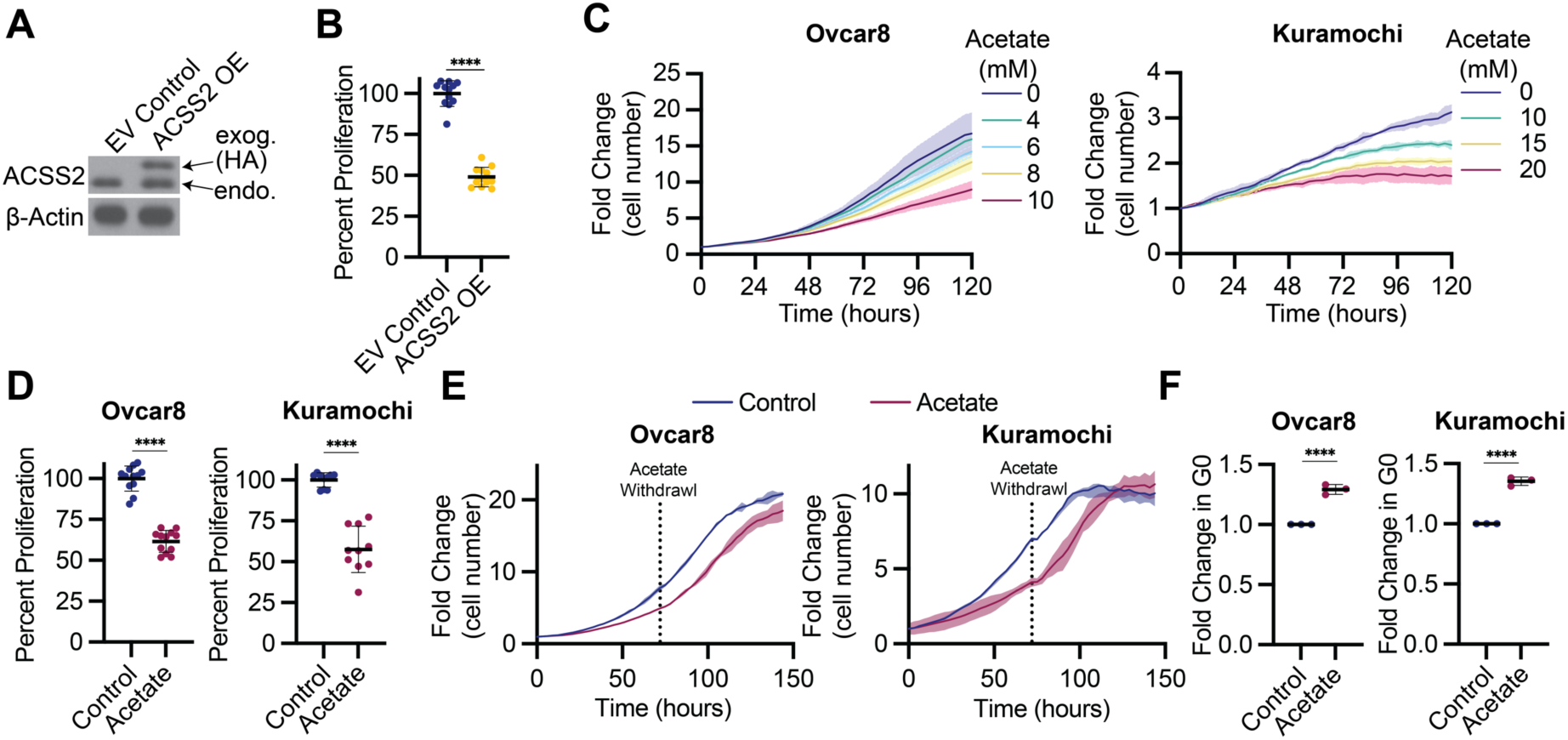
Acetate and ACSS2 drive quiescence in HGSC models. **(A)** Ovcar8 cells were transduced with lentivirus expressing empty vector control (EV control) or HA-tagged ACSS2 (ACSS2 OE). ACSS2 and HA protein expression was determined by western blotting. b-actin was used as a loading control. Immunoblots shown are representative data from at least 2 independent experiments. **(B)** Percent proliferation relative to EV control was assessed after 4 days. Cell counts were assessed using the Incucyte Live Cell Imaging system at endpoint. Pooled data from 3 independent biological replicates (n=4). Graphs represent mean ± SD. **(C)** Real time imaging of the proliferation of Ovcar8 and Kuramochi cells treated with the indicated doses of acetate. Cell numbers were normalized to the 0 h timepoint. Data represent 1 biological replicate (n=3) of at least 3 independent experiments. **(D)** Summary of endpoint proliferation data from (C). Pooled data from at least 3 independent experiments (n = 3-4). **(E)** Proliferation as assessed by real time imaging. Cell numbers were normalized to the 0 h timepoint. Data represent 1 biological replicate (n=3) of at least 3 independent experiments. Horizontal dotted line indicates when media was replaced with fresh media without acetate. **(F)** Ovcar8 and Kuramochi cells were transduced with a modified FUCCI reporter construct and treated with acetate. Quiescence (cells in G_0_) was assessed by flow cytometry. Shown is the CDT1/p27 double positive G_0_ population normalized to control. Pooled data from 3 independent experiments is shown. Graphs represent mean ± SD. ****p<0.001

**Figure 4.**
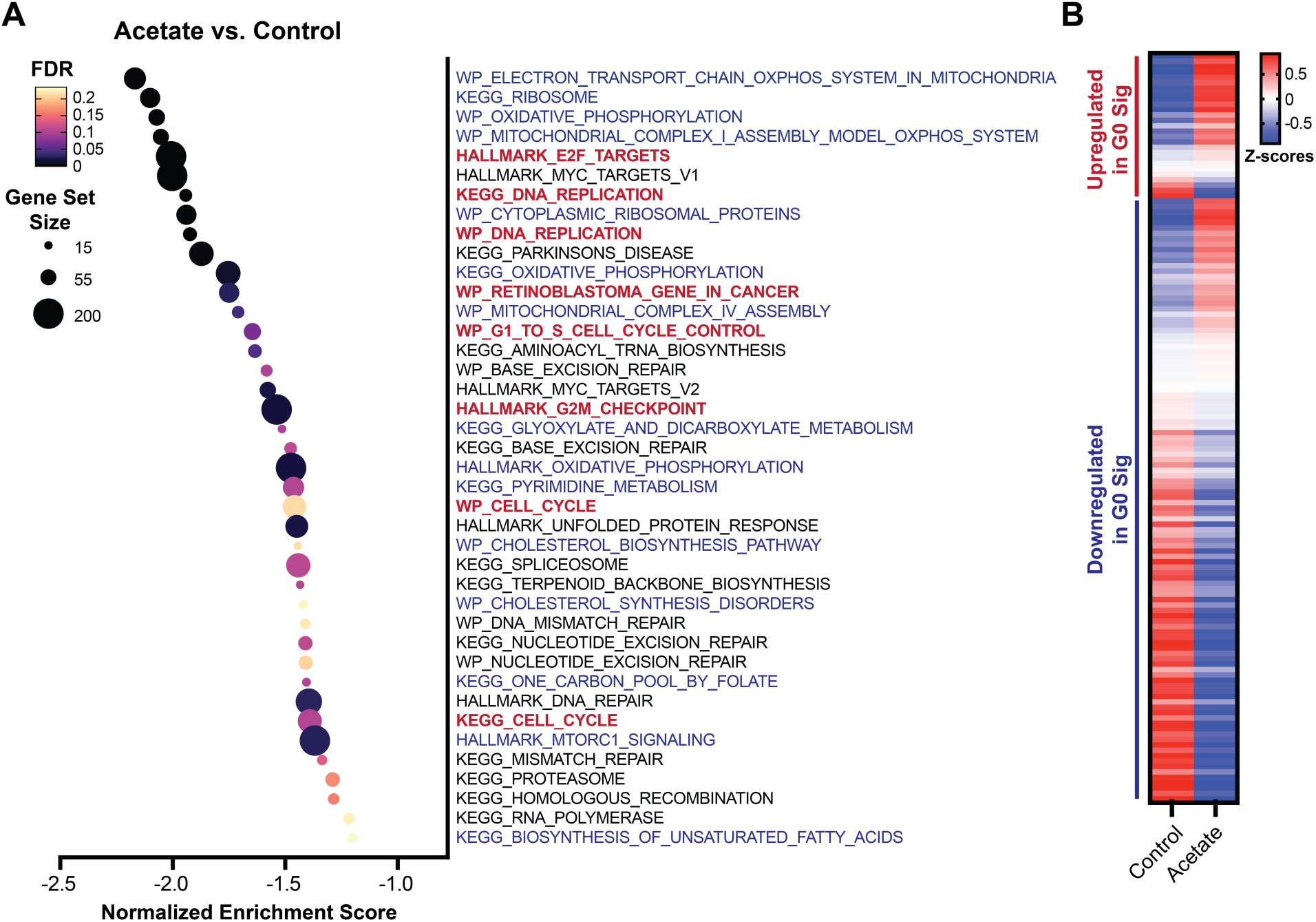
Acetate affects cell cycle transcriptional signatures in HGSC cells. **(A-B)** Ovcar8 cells were treated with 10mM acetate for 72h, and RNA-Seq was performed. **(A)** Negatively enriched Wikipathways (WP), Hallmark, and KEGG gene signatures in acetate vs. control. **(B)** Heatmap of genes in a published G0 signature [29].

### ACSS2 expression correlates with quiescence and poor outcome in patient samples

Finally, we evaluated the role of ACSS2 and quiescence in patient samples. High *ACSS2* expression correlates with worse both progression-free and overall survival in patients with ovarian serous carcinoma treated with platin (**Fig. 5A**). Consistent with the idea that increased ACSS2 is associated with quiescence, patients with a high quiescence gene signature [29] have moderately increased *ACSS2* expression (**Fig. 5B**). Moreover, high *ACSS2* expression corresponds with more platinum-resistant disease (**Fig. 5C**). As we observed increased ACSS2 in non-adherent conditions (**Fig. 1E**), we evaluated ACSS2 expression in matched solid tumor and ascites from HGSC patients. We found that ACSS2 expression is increased in cells from ascites compared to cells from the primary tumor (**Fig. 5D**). Together, these data demonstrate that ACSS2 expression is associated with quiescence and chemoresistance in clinical samples.

**Figure 5.**
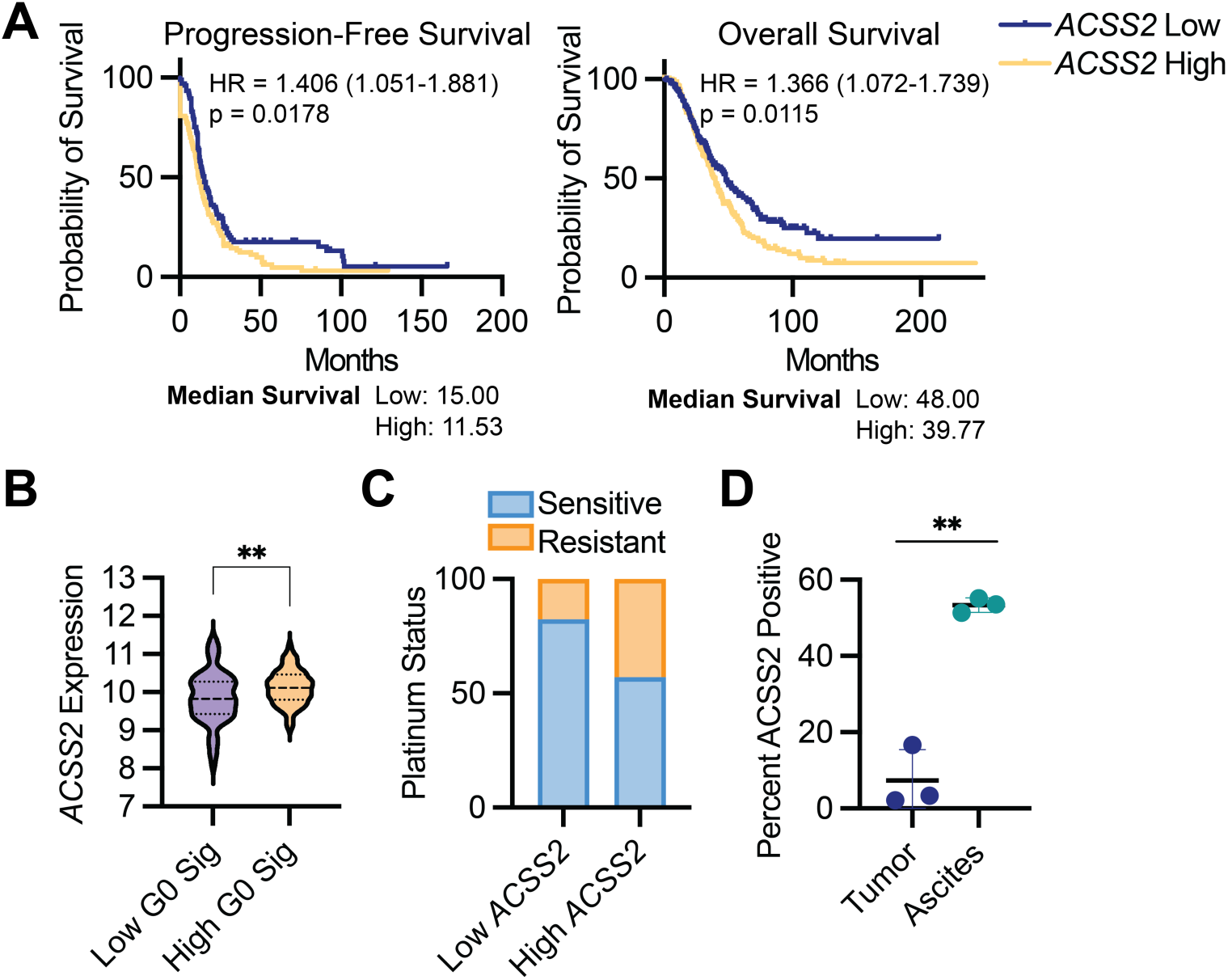
ACSS2 expression corresponds to worse outcomes and is increased in ascites compared to matched primary HGSC tumors. **(A)** Progression-free and overall survival in patients with ovarian serous carcinoma treated with platin stratified by *ACSS2* expression. Data from KMplot.com. **(B)** *ACSS2* expression in HGSC patients from TCGA stratified by their G0 signature [29]**. (C)** Platinum status in HGSC patients from TCGA with low and high *ACSS2* expression. **(D)** ACSS2 expression in matched primary tumor cells and ascites associated cells was determined by immunofluorescence. n = 3. Graphs represent mean ± SD. ****p<0.01

## Discussion

While quiescence cells were initially thought to be metabolically inactive [30], recent evidence has shed new light on metabolic plasticity of this cell state. In cancer, this is of paramount importance as quiescent cells contribute to chemoresistance. Whether metabolic reprogramming drives quiescence in ovarian cancer is underexplored. Here we identified an unexpected role for ACSS2-mediated acetyl-CoA metabolism in driving ovarian cancer quiescence.

Quiescence has been historically defined as a metabolically-inactive cell cycle exit [30]. Work in normal adult stem cells, which are quiescent, has indicated that quiescent cells may in fact rely on metabolic rewiring to retain their pluripotent or functional state [7, 8]. Similarly, metabolism is also important for quiescent stem cell survival, and prior reports have demonstrated a direct role for fatty acid oxidation (FAO) in this process in both normal and cancerous quiescent cells [7, 8, 31-33]. FAO is the breakdown of fatty acids into acetyl-CoA units [34]. Consistently, we found that quiescent cancer cells have increased acetyl-CoA (**Fig. 2B**), and we found that this acetyl-CoA is derived from acetate and likely not other sources (**Fig. 2C, 2E, and S3C**). Together, these data support an important role for acetyl-CoA-mediated metabolic reprogramming in supporting the quiescent cell state. Interestingly, senescence, another non-proliferative cell state, is well-known to be associated with metabolic reprogramming [35-41]. While further work is needed to understand the changes in metabolism in reversible versus irreversible cell cycle exits, we posit that metabolic reprogramming is a shared hallmark of non-proliferative cells.

How acetate and acetyl-CoA are promoting a quiescent state in our cells is an open question. Prior work demonstrates that acetyl-CoA is needed for histone acetylation that primes exit from quiescence [42-44]. In yeast, Acs2 (the yeast homolog of ACSS2) is recruited to chromatin during quiescence exit to participate in localized transfer of acetyl groups [42]. In cancer, ACSS2 and acetate support proliferation, growth, and survival under nutrient deprivation or hypoxia [45-50]. Mechanistically, studies have shown this is in part via nuclear ACSS2 localization [45]. In contrast, a recent paper reports that acetate limits cell proliferation in the presence of glucose, and high ACSS2 is associated with decreased tumor burden [51]. Indeed, our studies were performed in nutrient replete media. Thus, it is possible that both the nutrient status of cells and the subcellular localization of ACSS2 and acetate-mediated acetyl-CoA synthesis is critically important for either inducing or suppressing quiescence. Cytoplasmic acetate-derived acetyl-CoA via ACSS2 supports multiple metabolic processes, including cholesterol, fatty acid, isoprenoid, hexosamine, and ketone body synthesis in addition to providing acetyl groups for protein acetylation [52]. Future aims of our work are to determine which pathway is regulating quiescence induction in HGSC cells.

Chemoresistance is a major clinical issue for HGSC [10, 11]. Our prior work has supported the critical role for quiescent cells in HGSC chemoresistance [14, 15]. We have also demonstrated that metabolic reprogramming is associated with a decreased therapeutic response in HGSC [53]. Here we found that ACSS2 is upregulated in quiescent cells (**Fig. 1**), and high ACSS2 expression corresponds with worse outcomes in patients (**Fig. 5**).

Thus, it is interesting to speculate whether inhibiting ACSS2 will decrease quiescence and promote chemosensitization. Interestingly, prior studies have found that inhibition of ACSS2 decreases viability in spheroids [45], a model of quiescence, in addition to decreasing tumor burden *in vivo* [45, 54, 55]. This is thought to be in part due to ACSS2 upregulation under metabolic stress. Given our results that quiescent cells undergo a metabolic reprogramming (**Fig. 2**) and that ACSS2 expression corresponds to a G0 signature (**Fig. 5B**), it is possible that the observed effects in other studies are in part due to elimination of quiescent cells, although this will need to be tested. Nonetheless, given our results that increased *ACSS2* corresponds to platinum resistance in HGSC patient samples (**Fig. 5C**), targeting ACSS2 should be tested as an approach to delay or overcome chemoresistance in this patient population.

In summary, we provide evidence that acetate-derived acetyl-CoA metabolism drives HGSC quiescence and this axis is associated with worse patient outcomes. Future studies are needed to determine the mechanism of how ACSS2 and acetate drive quiescence in addition to the *in vitro* and *in vivo* effects of ACSS2 inhibitors on cisplatin response. Our data shed new light on the metabolic reprogramming that occurs in quiescent cancer cells.

## Acknowledgements

We would like to thank Fran Vazquez for help with the CRISPR schematic, Uma Chandran and Jiefei Wang (UPMC Hillman Cancer Center Bioinformatics Core) for help with bioinformatic analysis of the CRISPR screen, and Maureen Lyons (UPMC Hillman Cancer Center Cancer Genomics Core) for help with sequencing of the CRISPR library. This work was supported by National Institutes of Health (R37CA240625 to KMA, R01CA259111 to KMA and NWS, R01CA278100 and 5P50CA272218 to RJB), the American Cancer Society (RSG-19-113-01-CCG to KMA), a generous gift from M. Deitich, the Women’s Cancer Research Center Pilot Award (to ACS), and the UPMC Hillman Cancer Center. This project used the Cancer Genomics Facility and the Cancer Bioinformatics Services that are supported in part by award P30CA047904.

## Author Contributions

ACS: Investigation, Writing – Original Draft, Visualization, Supervision, Funding Acquisition. EM: Investigation, Formal Analysis. AJC: Investigation. AF: Investigation. SM: Investigation. NH: Conceptualization, Writing – Review & Editing. NWS: Conceptualization, Methodology, Investigation, Supervision, Writing – Review & Editing, Funding Acquisition. RJB: Conceptualization, Writing – Original Draft, Writing – Review & Editing, Supervision, Project Administration, Funding Acquisition. KMA: Conceptualization, Formal Analysis, Writing – Original Draft, Writing – Review & Editing, Visualization, Supervision, Project Administration, Funding Acquisition.

## Supplemental Figures

**Supplemental Figure 1.**
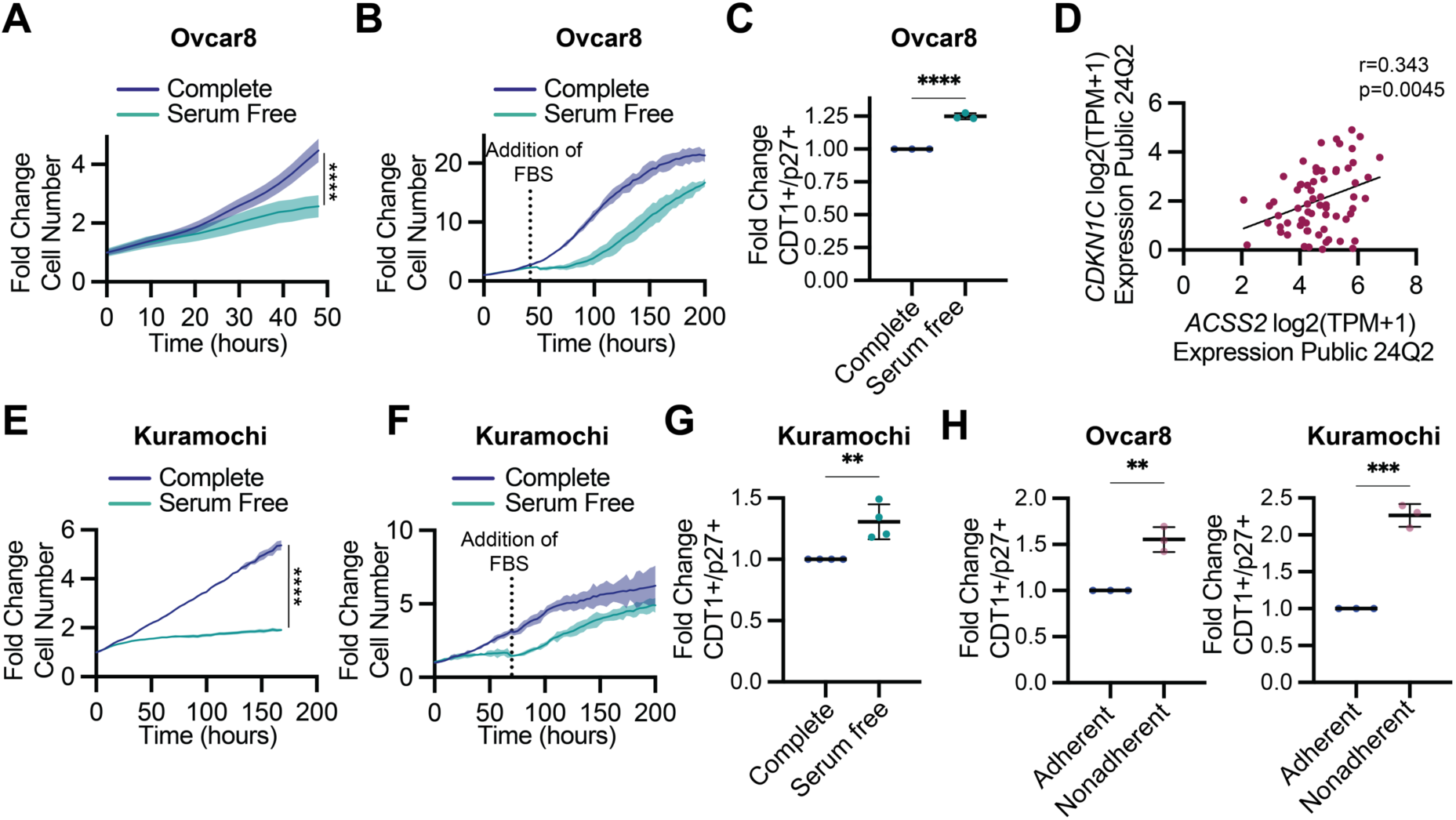
Induction of quiescence in HGSC cells by serum starvation and culturing in nonadherent conditions. Related to Figure 1. **(A)** Ovcar8 cells were cultured in complete or serum free media, and proliferation was assessed by live cell imaging. Data represent 1 of at least 2 independent experiments (n=3). (**B)** Same as (A), but the horizontal dotted line indicates when media was replaced with fresh media with 5% FBS. Data represent 1 biological replicate (n=3) of at least 2 independent experiments. **(C)** Ovcar8 cells were transduced with a modified FUCCI reporter construct and cultured in with or without serum. Quiescence (cells in G_0_) was assessed by flow cytometry. Shown is the CDT1/p27 double positive population normalized to control. Pooled data from 3 independent experiments. Graphs represent mean ± SD. **(D)** Pearson correlation between *ACSS2* and *CDKN1C* expression in ovarian/fallopian tube lineage cell lines in DepMap. **(E-G)** Same as (A-C) but in Kuramochi cells. **(H)** Cells were transduced with a modified FUCCI reporter construct and cultured on adherent or non-adherent conditions. Quiescence (cells in G_0_) was assessed by flow cytometry. Shown is the CDT1/p27 double positive population normalized to control. Data represent 3 independent experiments. Graphs represent mean ± SD. **p<0.01; ***p<0.005; ****p<0.001

**Supplemental Figure 2.**
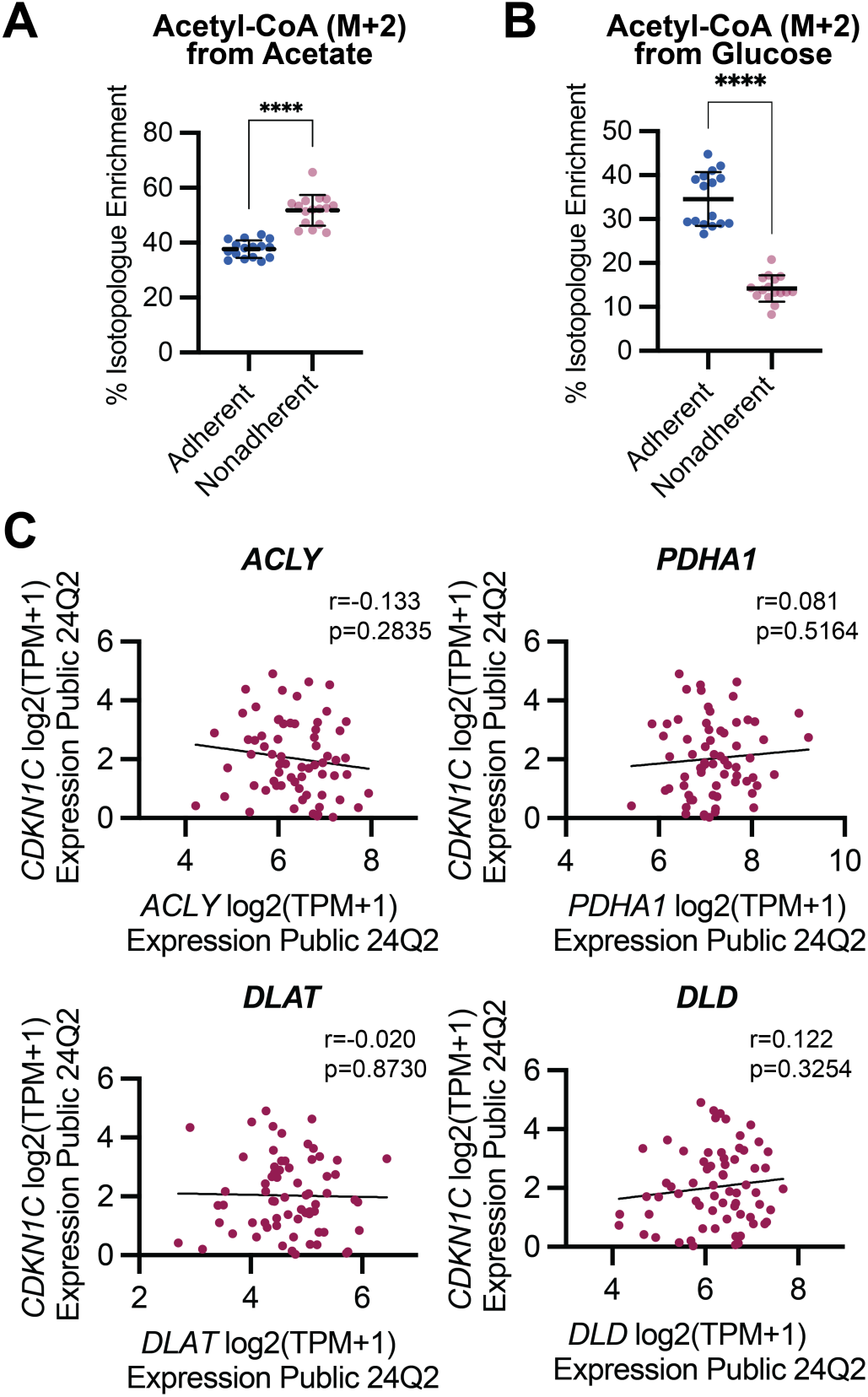
Nonadherent quiescent cells have increased acetate-derived acetyl-CoA; other acetyl-CoA producing enzymes do not correlate with *CDKN1C*. Related to Figure 2. **(A)** Acetyl-CoA M+2 percent isotopologue enrichment from labeled acetate in the indicated cells. Pooled data from 2 independent experiments (n=8). **(B)** Acetyl-CoA M+2 percent isotopologue enrichment from labeled glucose in the indicated cells. Pooled data from 2 independent experiments (n=8). **(C)** Pearson correlation between the indicated genes and *CDKN1C* expression in ovarian/fallopian tube lineage cell lines in DepMap. Graphs represent mean ± SD. ****p<0.001

**Supplemental Figure 3.**
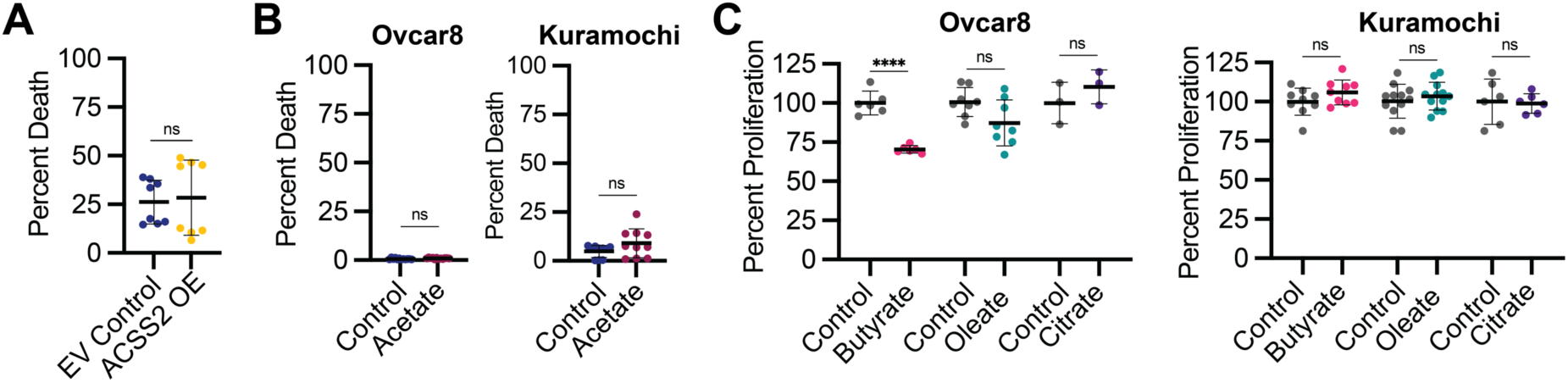
ACSS2 overexpression and acetate do not decrease viability; other acetyl-CoA sources do not decrease proliferation. Related to Figure 3. **(A)** Ovcar8 cells were transduced with lentivirus expressing empty vector control (EV control) or HA-tagged ACSS2 (ACSS2 OE). Percent death at endpoint (96 hrs) normalized to controls is shown. Pooled data from 2 independent experiments (n=4). **(B)** Ovcar8 and Kuramochi cells were supplemented with acetate. Percent death at endpoint is shown. Pooled data from at least 2 independent experiments (n=3). **(C)** Ovcar8 and Kuramochi cells were supplemented with sodium butyrate (Ovcar8: 100 mM, *n* = 2; Kuramochi: 40 mM, *n* = 3), oleate (Ovcar8: 200 mM, *n* = 3; Kuramochi: 100 mM, *n* = 4), or citrate (1 mM; Ovcar8: *n* = 1, Kuramochi: *n* = 2) for 72 hrs. Pooled percent cell number at endpoint normalized to controls is shown. Graphs represent mean ± SD. ****p<0.001; ns = not significant

